# Expression of the cyanobacterial F_0_F_1_ ATP synthase inhibitor AtpΘ depends on small basic DNA-binding proteins and differential mRNA stability

**DOI:** 10.1101/2021.11.17.468837

**Authors:** Kuo Song, Martin Hagemann, Jens Georg, Sandra Maaß, Dörte Becher, Wolfgang R. Hess

## Abstract

F_0_F_1_ ATP synthases produce ATP, the universal biological energy source. ATP synthase complexes on cyanobacterial thylakoid membranes use proton gradients generated either by photosynthesis or respiration. AtpΘ is an ATP synthase regulator in cyanobacteria which is encoded by the gene *atpT*. AtpΘ inhibits the hydrolysis of ATP (reverse reaction) that otherwise would occur under unfavorable conditions. In the cyanobacterium *Synechocystis* sp. PCC 6803, AtpΘ is expressed maximum in darkness but at very low levels under optimum phototrophic growth conditions or in the presence of glucose. DNA coimmunoprecipitation experiments followed by mass spectrometry identified the binding of the two transcriptional regulators cyAbrB1 and cyAbrB2 to the promoter and the histone-like protein HU to the 5’UTR of *atpT*. Analyses of nucleotide substitutions in the promoter and GFP reporter assays identified a functionally relevant sequence motif resembling the HLR1 element bound by the RpaB transcription factor. Electrophoretic mobility shift assays confirmed interaction of cyAbrB1, cyAbrB2 and RpaB with the promoter DNA. However, overall the effect of transcriptional regulation was comparatively low. In contrast, *atpT* transcript stabilities differed dramatically, half-lives were 1.6 min in the light, 33 min in the dark and substantial changes were observed if glucose or DCMU were added. These findings show that basic transcriptional control of *atpT* involves nucleoid-associated DNA-binding proteins, positive regulation through RpaB, while the major effect on the condition-dependent regulation of *atpT* expression is mediated by controlling mRNA stability, which is related to the cellular redox and energy status.

**IMPORTANCE:** F_0_F_1_ ATP synthases are protein complexes that produce ATP, the universal biological energy source in all kinds of organisms. Under unfavorable conditions, ATP synthases can operate in a futile reverse reaction, pumping protons while ATP is used up. Cyanobacteria perform plant-like photosynthesis but they cannot use the same mechanism as plants that inhibit chloroplast ATP synthases entirely during the night because respiratory and photosynthetic complexes are both located in the same membrane system. AtpΘ is a small peptide inhibitor of the reverse ATPase function in cyanobacteria encoded by the gene *atpT*. The production of AtpΘ is highly regulated to ensure that it is only synthetized when it is needed. In the here presented work we found that three transcription factors contribute to the regulation of *atpT* expression. However, we identified the control of mRNA stability as the major regulatory process governing *atpT* expression. Thus, it is the interplay between transcriptional and posttranscriptional regulation that position the AtpΘ-based inhibitory mechanism within the context of the cellular redox and energy balance.

## Introduction

F_0_F_1_ ATP synthase (hereafter referred to as ATP synthase) is a multisubunit enzyme complex that resides in the energy-transducing membranes of chloroplasts, mitochondria, bacteria and archaea (1). ATP synthases from different organisms or organelles share a similar architecture that consists of a membrane-extrinsic F_1_ head and a membrane-intrinsic F_0_ complex connected by central and peripheral stalks (2). The catalytic activity resides in the F_1_ part of the complex, which synthesizes ATP from ADP and P_i_ using proton motive force (Δ*p*). Under unfavorable conditions, ATP synthases operates backward, hydrolyzing ATP while pumping protons. Therefore, multiple mechanisms have evolved to inhibit this futile ATP hydrolysis activity *in vivo*.

Small peptide inhibitors were identified for ATP synthases in mitochondria. They are known as inhibitory factor 1 (IF1) in mammals (3, 4), while three inhibitors, IF1, STF1 and STF2, have been described in yeast (5, 6). IF1 inhibits the ATPase activity of mitochondrial ATP synthase under depolarizing conditions such as hypoxia (2, 7, 8). A different mechanism inhibits the reverse reaction of ATP synthase in chloroplasts, where the proton gradient is generated by light-driven photosynthetic electron transport. In this plant organelle, a redox-controlled mechanism switches ATP synthase activity off at night when photosynthesis does not occur (9). The molecular mechanism underlying this redox switch relies on the gamma subunit of chloroplast ATP synthases, which forms a disulfide bond through cysteine residues in the unique sequence motif EICDINGXC, inhibiting ATP hydrolysis activity (9–11).

Cyanobacteria differ uniquely from all other bacteria and eukaryotic energy-converting organelles by combining photosynthetic and respiratory electron transfer systems in their thylakoid membranes. Consequently, two fundamentally distinct systems generate the proton gradient utilized by ATP synthase. Therefore, regulatory mechanisms are expected for the ATP synthase of cyanobacteria that resemble those of mitochondria and chloroplasts, as well as unique mechanisms. Although photosynthetic and respiratory electron transfer systems in cyanobacteria are interconnected by multiple possible electron transport routes (12), the mechanisms controlling the different electron transport routes during fluctuating conditions are largely unclear (13).

Indeed, cyanobacteria encode a small amphipathic protein, AtpΘ, which was characterized as an inhibitor of ATP synthase ATPase activity (14). AtpΘ is encoded by the *atpT* gene, and likely homologs of *atpT* were detected in all available cyanobacterial genomes. In the model cyanobacterium *Synechocystis* sp. PCC 6803 (*Synechocystis* 6803), AtpΘ accumulates during conditions that potentially weaken the proton gradient, such as darkness (14, 15). The addition of glucose neutralized the strong dark-induced *atpT* transcript accumulation, while the further addition of a chemical uncoupler or an electron transport inhibitor restored the high transcript accumulation in the dark (14). These findings suggested that the regulation of *atpT* is complex and to be somehow connected to the status of the electron transport chain or proton gradient.

The present study investigated mechanisms involved in regulating *atpT* expression. We first analyzed the *Synechocystis* 6803 *atpT* promoter by systematically mutating the region upstream of the transcriptional start site (TSS) in a GFP-based reporter assay. Transcript accumulation and promoter activity were then investigated under different conditions by combining the GFP assay with analyses of mRNA accumulation and stability. Protein coimmunoprecipitation experiments using a biotinylated *atpT* promoter fragment identified cyAbrB1 and cyAbrB2 as interacting transcription factors. The interaction of these factors as well as of a third transcription factor, RpaB, was confirmed in electrophoretic mobility shift assays *in vitro*. However, we found that the major level of control occurs at the post—transcriptional level. The data provide insights into the complex regulation of *atpT* expression and facilitate further studies of the underlying regulatory mechanisms and AtpΘ functions.

## Results

### The *atpT* promoter contains two regions for basal activity and inducibility by darkness

The TSS of *atpT* is located at nt position 299115 on the reverse strand of the *Synechocystis* 6803 chromosome (GenBank file NC_000911), yielding a 5’UTR of 143 nt, a coding region of 144 nt, and a 3’UTR of 101 nt (Fig. S2A). The 3’UTR ends with a predicted stem-loop structure that resembles a Rho-independent terminator (Fig. S2B). According to a previous study, the *atpT* (called *norf1* in that study) transcript is strongly regulated and becomes the most abundant mRNA after 12 h in the dark (25). Because the location of regulatory sequence elements was unknown, a 328 nt fragment upstream of the TSS was empirically chosen (P_*atpT328*_). Based on the detected sequence conservation in the TSS-proximal promoter region among cyanobacteria (Fig. S3), another two shorter versions (P_*atpT94*_ and P_*atpT66*_) were also selected for our experimental analyses.

A GFP reporter gene assay was performed to study the promoter activity *in vivo*. As indicated in Fig. 1A, the selected promoter fragments were combined with different 5’UTRs followed by an sf*gfp* cassette, fused into the pVZ322 backbone and conjugated into *Synechocystis* 6803. Constructs used for GFP assays comprised one of the three *atpT* promoter variants or the constitutive BBa_J23101 promoter (26) and the native *atpT* 5’UTR or the 29 nt 5’UTR of housekeeping gene *sll0639* (Fig. 1B). The *sll0639* 5’UTR belongs to a transcript that is relatively abundant and stably accumulated (25). The GFP intensities of the four strains containing the full-length promoters were measured over 48 h. At the time point of 0 h, the cells were transferred to the dark, followed by a return to the light at 24 h. The combination of P_*atpT328*_ and native 5’UTR yielded increasing GFP intensities after transfer to the dark and decreased GFP intensities during the subsequent light period. Furthermore, this promoter yielded the highest intensities at all time points by far, including the light condition (Fig. 1C, GFPconstruct_1). Replacement of the native *atpT* 5’UTR with the *sll0639* 5’UTR led to considerably lower GFP intensity. If transcription was driven by the BBa_J23101 promoter, fluorescence was lower for the respective 5’UTR-sf*gfp* fusions than those controlled by P_*atpT328*_. However, again, fluorescence was higher if the native 5’UTR was used rather than the *sll0639* 5’UTR. We conclude that the *atpT* promoter is comparatively strong and that the native 5’UTR contributes to the observed high expression of the *atpT* gene.

**FIG 1.**
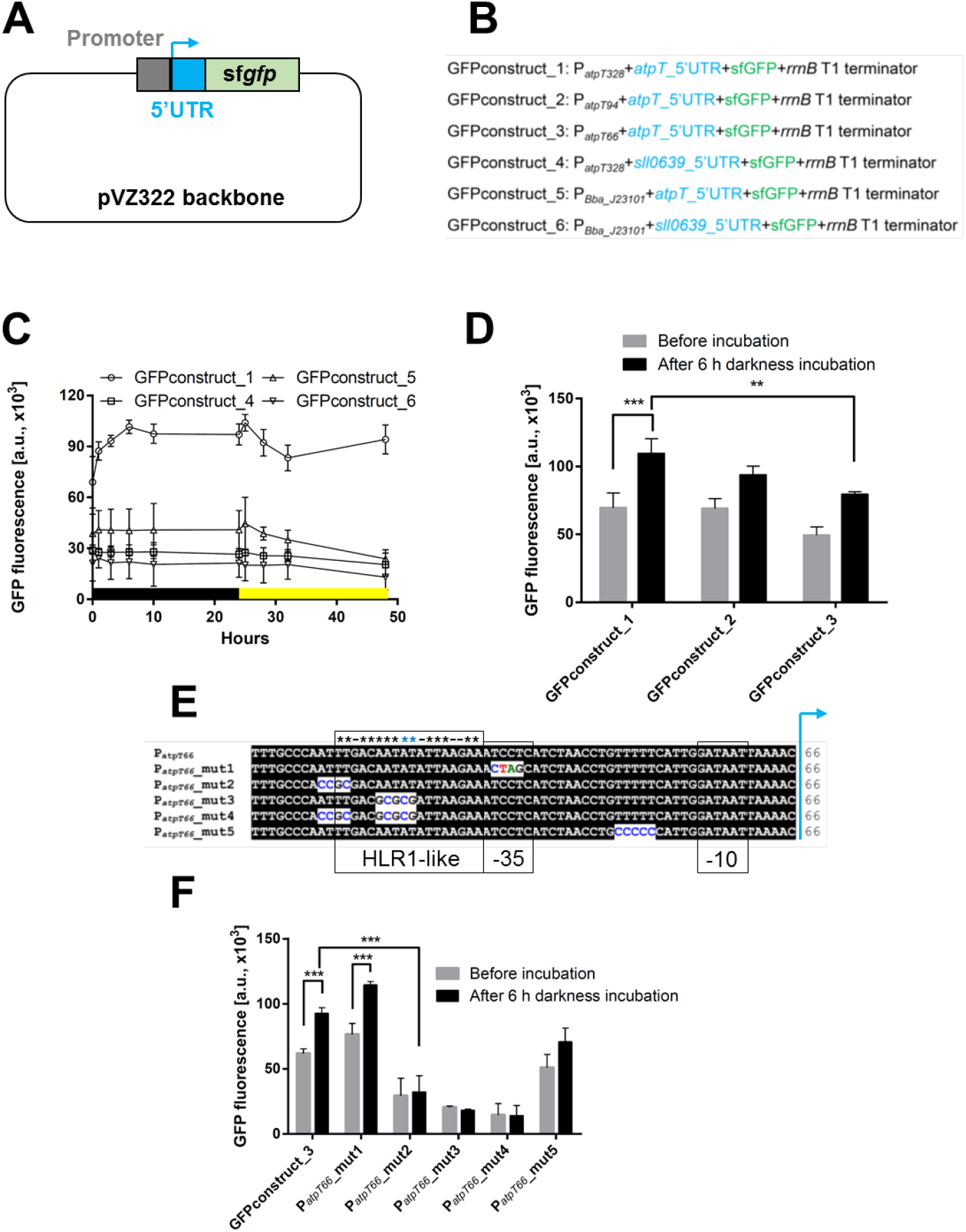
Reporter gene assay to monitor transcription driven by different promoter variants. (A) General design of the plasmids used for the GFP assay. Using pVZ322 as the backbone, the coding sequence of sf*gfp* together with the *rrnB* T1 terminator sequence (47) was fused to various 5’UTR (blue) and promoter (gray) sequences. (B) The design of different constructs used for the GFP assays. In constructs 1, 4, 5 and 6, two different 5’UTRs were combined with the full-length *atpT* promoter P_*atpT328*_ or the P_*Bba_J23101*_ promoter ((iGEM Registry parts.igem.org (26)). Constructs 2 and 3 contain truncated versions of the *atpT* promoter. The 5’UTRs originated from *atpT* or the *sll0639* gene encoding the oligoribonuclease/PAP phosphatase NrnA. (C) Analysis of the GFP intensity in dark/light shift experiments. Different strains cultured under continuous light were first transferred to darkness for 24 h (black bar) and then transferred to light (yellow bar). Signals measured from a simultaneously cultivated *Synechocystis* 6803 without any genetic modification were subtracted from the measured fluorescence values (background correction). (D) GFP assay comparing the activities of *atpT* promoters of three different lengths before (gray bars) and after 6 h of dark incubation (black bars). (E) Mutations introduced into the P_*atpT66*_ promoter to test the relevance of conserved sequence elements. The mutated nucleotides, putative -35 and -10 elements and the region with similarity to the HLR1 element (two direct repeats (G/T)TTACA(T/A)(T/A) separated by 2 nt (20)) is boxed and asterisks indicate matching positions. (F) GFP assay comparing the activities of P_*atpT66*_-5’UTR and the five of its mutated variants as depicted in Panel (E) before (gray bars) and after 6 h of dark incubation (black bars). The differences between groups were analyzed using *t* tests (Panels (D) and (F); Tables S5 and S6) with GraphPad software. Significance was established at *P* < 0.05 = ** and *P* < 0.01 = ***. For details of the statistical analysis see Tables S5 and S6.

After transfer to darkness, an increased GFP intensity was only observed for the GFPconstruct_1 in which the *atpT* promoter and native 5’UTR were combined, while the replacement of either the 5’UTR (GFPconstruct_4) or the promoter (GFPconstruct_5) led to the loss of dark induction. Based on these results, the combination of the native promoter and 5’UTR is required for increased expression in the dark.

Based on the time course of changes in GFP intensity displayed in Fig. 1C, 6 h of dark incubation were used in subsequent assays to analyze the promoter activity. In this experiment, the activities of three *atpT* promoters of different lengths fused to the *atpT* 5’UTR were measured (Fig. 1D). Compared with P_*atpT328*_, the promoter P_*atpT66*_ showed significantly lower activity under both light and dark conditions, while P_*atpT94*_ had similar activity under light conditions but lower activity under dark conditions. Therefore, the *atpT* promoter is regulated at different positions and probably by different regulators.

The P_*atpT66*_ promoter was mutated at three sites separately or in combination, as shown in Fig. 1E, to study the functional relevance of conserved nucleotides, which were identified in promoter sequence comparisons of *atpT* genes from several cyanobacteria (Fig. S3). Among these mutations, P_*atpT66*__mut1 contains a substitution of 4 nt within the putative -35 region, P_*atpT66*__mut2 and P_*atpT66*__mut3 contain tetra- and pentanucleotide substitutions of two AT-rich regions, respectively, while in P_*atpT66*__mut4, both AT-rich regions were replaced. In construct P_*atpT66*__mut5, a stretch of five consecutive thymidine residues from position -18 to -23 was replaced with cytosine nucleotides.

The activities of these promoters were elucidated with the GFP assay. As shown in Fig. 1F, P_*atpT66*__mut2-, P_*atpT66*__mut3- and P_*atpT66*__mut4-containing mutations in the AT-rich segments showed no promoter activity, indicating that this region is essential for the transcription of the downstream gene. In contrast, mutation of the putative -35 region in P_*atpT66*__mut1 had no discernible effect, consistent with previous findings that the majority of promoters in *Synechocystis* 6803 lacks a -35 element (27). The construct P_*atpT66*__mut5 showed a somewhat lower GFP intensity than P_*atpT66*_, indicating that this segment was of minor relevance. However, all of the active promoters showed higher activities in the dark than in light, similar to the wild-type promoter. We conclude that the activity of the *atpT* promoter is independent of a -35 element, but that the two AT-rich regions from positions -58 to -55 and position -51 to -48 are crucial for its activity.

### Effects of inhibitors on *atpT* promoter activity and transcript accumulation

In *Synechocystis* 6803, the *atpT* transcript was detectable under all tested conditions (25), but its level increased substantially under low-energy conditions such as darkness (15). We studied transcript accumulation in the presence of glucose or different inhibitors to discern between different, possibly relevant effectors. The *atpT* mRNA level was low but detectable at a light intensity of 40 µmol photons m^-2^ s^-1^ but much higher in the dark, and the addition of 10 mM glucose neutralized the increased mRNA accumulation in darkness (Fig. 2A and B), consistent with previous results (14). This fact excludes the possibility that the dark induction of *atpT* expression was controlled by a photoreceptor. The addition of DCMU led to a higher *atpT* mRNA level, while the presence of the uncoupler carbonyl cyanide m-chlorophenyl hydrazone (CCCP) (28) or methylviologen had no discernible effects on *atpT* levels (Fig. 2A and B).

**FIG 2.**
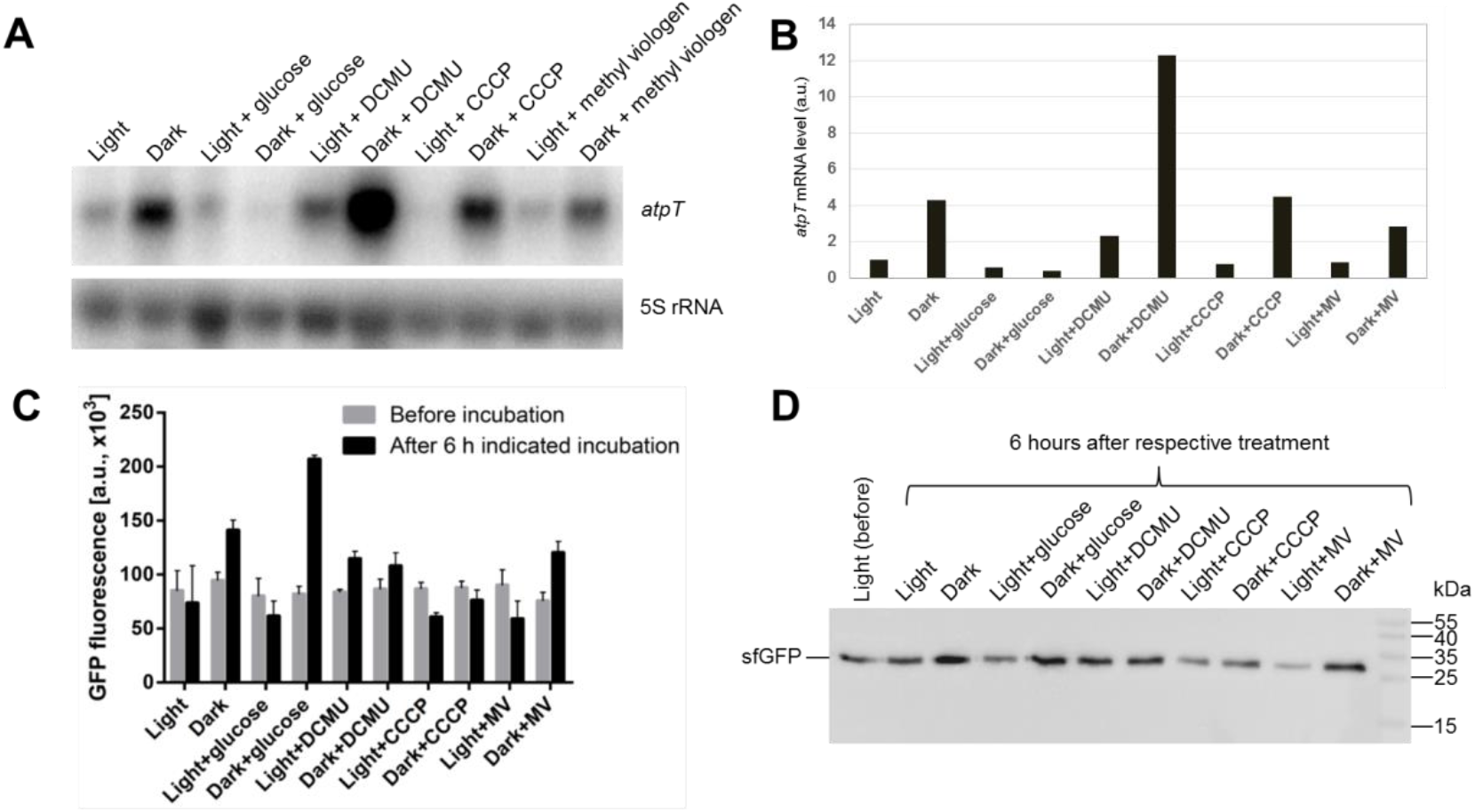
*atpT* transcript accumulation and reporter gene assays under different conditions. (A) Northern blot analysis of *atpT* mRNA accumulation in the wild type strain. Total RNA samples were isolated from exponential phase cultures incubated under the indicated conditions for 6 h each. 5S rRNA was used as a loading control. One representative blot is shown (n=2). (B) Quantification of *atpT* transcript levels shown in Panel (A) after normalization to 5S rRNA levels. (C) GFP reporter gene assay comparing *atpT* promoter activity before (gray bars) and after 6 h of incubation under the indicated conditions (black bars). Exponential phase cultures were first adjusted to the same OD750 value and cultured overnight, and then the GFP intensity was measured to acquire the intensities before incubation and after 6 h of treatment. (D) Representative immunoblot analysis of the expression of GFP in strains cultivated under the same conditions as described in Panel (C) (n=2). The same amounts of total protein (20 μg) isolated from *Synechocystis* 6803 cells before treatment (light) or after 6 h of the indicated treatment were loaded. Signals were detected with a rabbit anti-sfGFP antibody (1:2000 dilution) followed by anti-rabbit IgG secondary antibody (1:5000 dilution). PageRuler Prestained Protein Ladder (10–170 kDa, Fermentas) was used as a molecular mass marker.

The observed effects on steady-state *atpT* mRNA levels might result from altered transcription (promoter activity), transcript stability, or both. GFP reporter gene assays were conducted using the strain bearing GFPconstruct_1 under the same conditions as shown in Fig. 2A to distinguish between these possibilities. An incubation in the dark for 6 h led to a higher GFP intensity (Fig. 2C), consistent with the results of reporter gene assays shown in Fig. 1C and transcript accumulation data shown in Fig. 2A and B. DCMU increased the GFP intensity under light and dark conditions (consistent with the RNA data shown in Fig. 2A and B), while CCCP induced weak dampening and methylviologen exerted a weak inverse effect under both conditions. An unexpected result was observed if glucose was added. In the presence of 10 mM glucose, the GFP intensity in the dark increased more substantially than that observed in the dark alone. A western blot was performed to verify this result and showed maximum accumulation of sfGFP in the dark and in darkness plus glucose (Fig. 2D). Hence, this effect of glucose on the expression of the reporter gene was in striking contrast to the Northern blot data in Fig. 2A, in which the addition of glucose during the dark incubation led to substantially diminished *atpT* mRNA accumulation. The effects of glucose stimulation in the dark on GFP fluorescence and protein abundance suggests that the dark-stimulated *atpT* promoter became even more active when glucose was added, while the amounts of *atpT* mRNA (but not the sf*gfp* mRNA) decreased, indicating a very strong effect of *atpT*-specific posttranscriptional RNA degradation.

### The *atpT* transcript exhibits conditionally distinct stabilities

Because the steady-state transcript level depends on both transcript synthesis and degradation, i.e., transcript turnover rates, the stabilities of the *atpT* mRNA were analyzed under different conditions, and their intensities relative to the 5S rRNA were calculated (Fig. 3). In the dark, the *atpT* transcript was degraded at a relatively slow decay kinetics, yielding an effective half-life of approximately 33 min (Fig. 3A and F). In the light, the transcript signal intensity deteriorated rapidly with a half-life of approximately 1.5 min (Fig. 3B and F). If glucose was added in the dark, transcript degradation was accelerated again, yielding a half-life of approximately 13 min (Fig. 3C and F).

**FIG 3.**
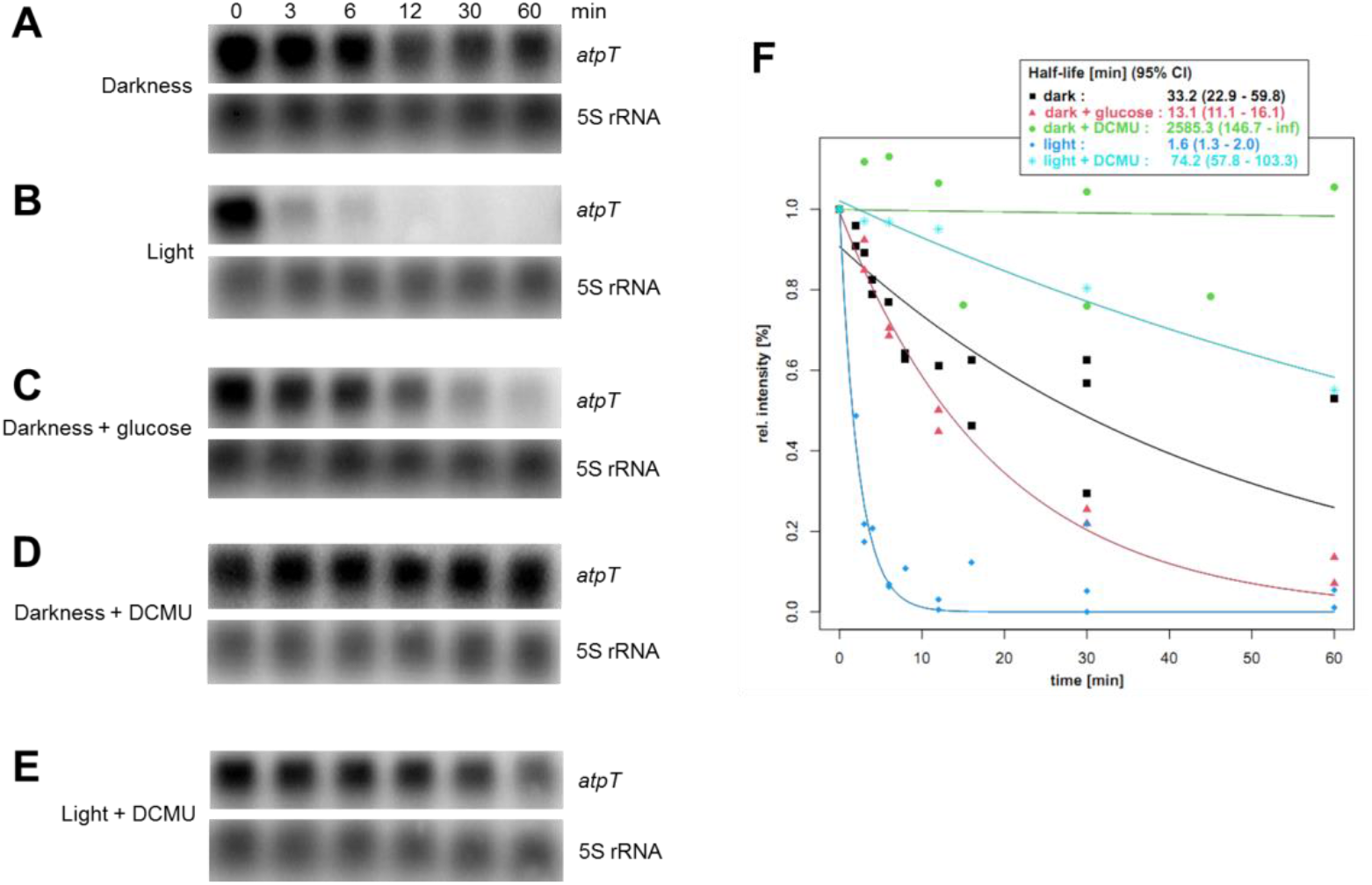
Stability of *atpT* transcripts under different conditions. (A-E) Levels of *atpT* mRNA detected in wild-type *Synechocystis* 6803 using Northern blot hybridization. The cultures were incubated in the dark overnight to induce high *atpT* mRNA levels, rifampicin was added, and the cultivation continued under the indicated conditions for 60 min. 5S rRNA was used as a loading control. Note that different exposure times were used for the different experiments. (F) Signal intensities of *atpT* from Panels (A-E) normalized for the respective 5S rRNA intensity plotted against the time after rifampicin addition. Quantity One software was used to quantify transcript signal intensities. The fitted decay curves are plotted as lines colored according to the respective conditions. Numerical values for half-lives in minutes and 95% confidence intervals (95% CI) are also provided. All stability analyses were repeated at least twice.

The addition of DCMU stabilized the *atpT* transcript under both light and dark conditions (Fig. 3D to F). This effect was not due to a nonspecific effect of DCMU on transcript stability because the level of the 5S rRNA hybridized on the same membranes did not show differences (Fig. 3A to E)., Under darkness + DCMU condition, the transcript was effectively stable, i.e., it showed no degradation during the measured period (Fig. 3F). These results suggest that different mRNA stabilities contributed very significantly to the observed differences in *atpT* expression levels under dark and light conditions.

### Identification of proteins interacting with the *atpT* promoter

Proteins binding to the *atpT* promoter were searched using a biotinylated P_*atpT66*_-5’UTR DNA fragment with the combination of pull-down and mass spectrometry to identify possible transcriptional regulators. In parallel, a DNA fragment containing only the biotinylated 5’UTR (without the upstream promoter sequence) was used. The bait fragments were incubated with soluble fractions of the whole cell lysate isolated from both light and dark cultures. The interacting proteins were first separated on SDS– PAGE gel (Fig. S4). While no qualitative difference was observed between proteins extracted from the light or dark samples, SDS–PAGE showed obvious differences between the proteins enriched by binding to the P_*atpT66*_-5’UTR or the 5’UTR fragments (indicated by arrows in Fig. S4). The proteins were then identified by mass spectrometry. The detailed mass spectrometry results are listed in Table S3.

The most notable differences observed on the SDS–PAGE gels were two abundant protein bands with apparent molecular masses of approximately 14 and 13 kDa in the P_*atpT66*_-5’UTR sample and a somewhat smaller protein band enriched with the 5’UTR fragment (Fig. S4). The mass spectrometry analysis identified the proteins cyAbrB1 and cyAbrB2 (Sll0359 and Sll0822 (16, 29)). Hence, two particular DNA-binding proteins were enriched in this size class (theoretical masses of 17.22 and 13.96 kDa) in the P_*atpT66*_-5’UTR sample, while the histone-like DNA-binding protein HU (30) Sll1712 (calculated molecular mass of 13.49 kDa) was identified as enriched by binding to the 5’UTR fragment (Table S3). Several additional proteins known to interact with nucleic acids were obtained from both samples in approximately similar quantities. These proteins were the transcription factor LexA, the DNA gyrase subunits A and B, and the ribonucleases PNPase, RNase E, RNAse J and RNase II (Table S3). The candidates for the other differentially enriched proteins observed on the SDS–PAGE gel (Fig. S4) located at approximately 25 kDa and 35 kDa might correspond to the Slr0637/Sll1471/Slr2051 and Sll1317/Slr1963/Sll0576 proteins. Among these candidates, Slr0637 is a hypothetical protein with unknown functions; Sll1471 and Slr2051 are two phycobilisome rod-core linker polypeptides that are components of the phycobilisome; Sll1317 is the Cytf subunit of the Cytb_6_f complex; Slr1963 is the orange carotenoid protein (OCP), which has a photoprotective role (31); and Sll1317 is a putative sugar-nucleotide epimerase/dehydratase. Due to their unlikely functions as specific transcriptional regulators, these proteins were not considered further.

However, the cyanobacterial AbrB-like transcriptional regulators (cyAbrBs) Sll0359 and Sll0822 have previously been reported to play important physiological roles in *Synechocystis* 6803. In particular, they function in carbon and nitrogen metabolism and, interestingly, in regulating the switch between day and night metabolic functions (16, 29, 32–35). Therefore, the roles of cyAbrB1 and cyAbrB2 in the regulation of *atpT* were further investigated.

### cyAbrB1 and cyAbrB2 interact with the *atpT* promoter *in vitro*

An N-terminal 6*His-tag was fused to both proteins, and the recombinant proteins were expressed in *E. coli* BL21 (DE3) and purified using affinity chromatography to test the binding of cyAbrB1 and cyAbrB2 to the *atpT* promoter. The purity of the obtained recombinant proteins was confirmed using SDS–PAGE (Fig. S5A). Then, electrophoretic mobility shift assays were performed to test their capacities for interacting with the *atpT* promoter and its 5’UTR. Both proteins showed specific interactions with the promoter when added at rather low levels, while in the presence of higher amounts (16 μg of protein for cyAbrB1 and 2.2 μg of AbrB2), the interaction with the 5’UTR DNA fragment was also observed (Fig. 4A and B). The hypershift (band at a higher position in Fig. 4B) observed when 16 μg of cyAbrB1 were applied might be explained by the abundant protein dimer observed on native SDS–PAGE gels (NSDS–PAGE; Fig. S5B). A dimeric form was also previously reported for cyAbrB2 *in vivo* (16), while the results in Fig. S5B suggested that cyAbrB2 was mostly monomeric *in vitro* under our conditions.

**FIG 4.**
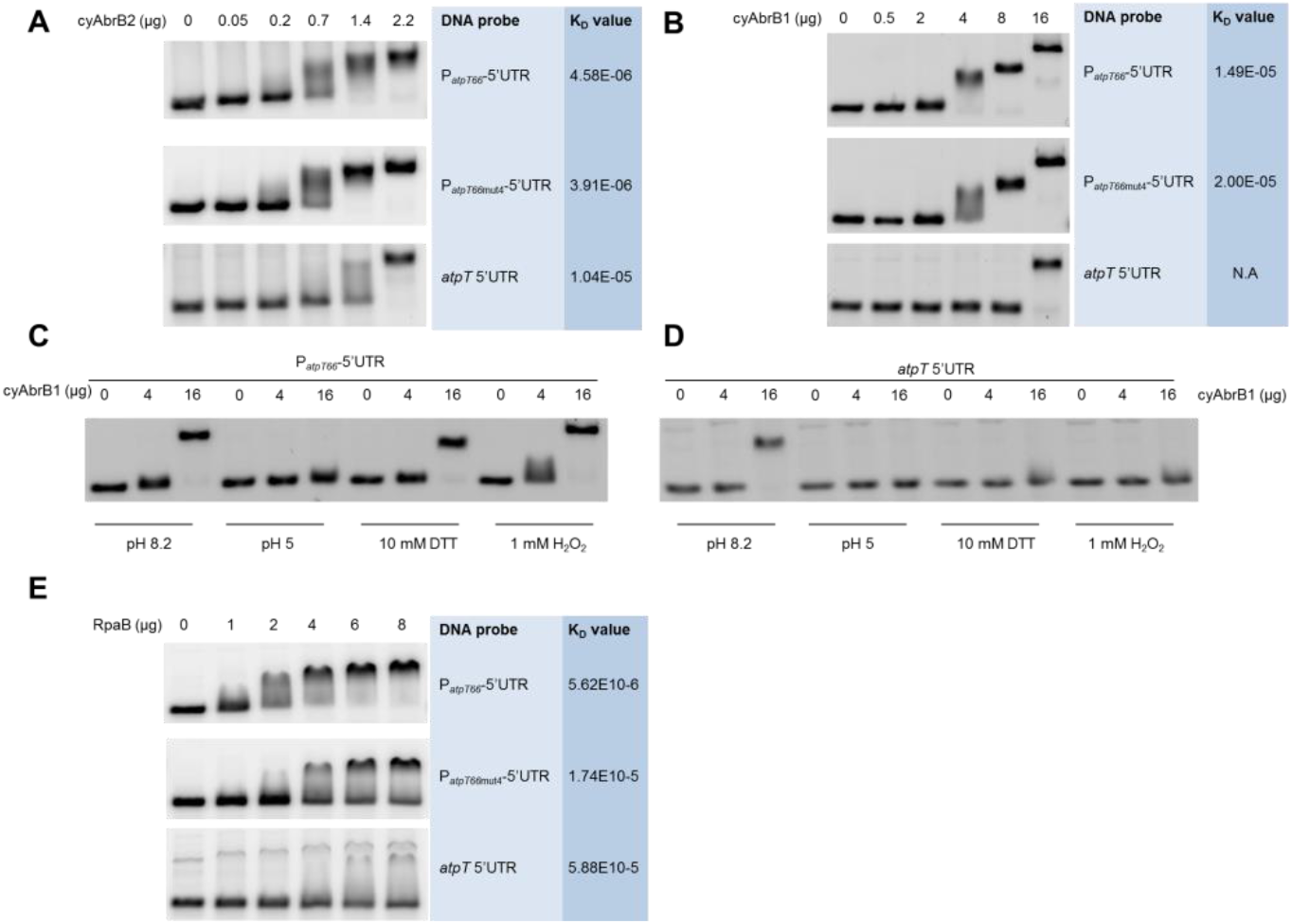
Electrophoretic mobility shift assays. (A) Binding of cyAbrB2 to three different DNA fragments. An increasing amount of purified His-tagged Sll0822 was incubated with 3.85 pmol of P_*atpT66*_-5’UTR, P_*atpT66mut4*_-5’UTR or *atpT* 5’UTR fragments labeled with Cy3 and then loaded onto 3% agarose gels for separation. The difference between P_*atpT66mut4*_-5’UTR and the native sequence is shown in Fig. 1E. (B) Binding of cyAbrB1 to the same DNA fragments as in Panel (A) but using only 0.385 pmol of each fragment. (C) Interaction of cyAbrB1 with the P_*atpT66*_-5’UTR DNA fragment at different pH values in the presence of DTT or H_2_O_2_. (D) Interaction of cyAbrB1 with the *atpT* 5’UTR fragment under the same conditions as described in Panel (C). (E) Binding of RpaB to the three DNA fragments as in (A) and (B). The calculated K_D_ values are listed next to the respective protein-DNA pairs. The K_D_ value between cyAbrB1 and a*tpT* 5’UTR could not be calculated (N.A), since no lane showing the intermediate state of binding was available. The sequences of DNA fragments used for gel shift assays are given in Table S4.

The binding of both cyAbrBs was undisturbed by the replacement of four and five conserved AT residues in the P_*atpT*_ promoter (construct P_*atpT66*mut4_-5’UTR, cf. Fig. 1E) because patterns identical to those of the P_*atpT66*_-5’UTR fragment were obtained, and the calculated K_D_ values showed no significant difference (Fig. 5A and B). Hence, these mutations impaired the *in vivo* promoter activity, while they had no effects on the *in vitro* binding of the cyAbrBs to the promoter region. We conclude that the AT-rich region must operate independently from binding to cyAbrB1 or cyAbrB2.

**FIG 5.**
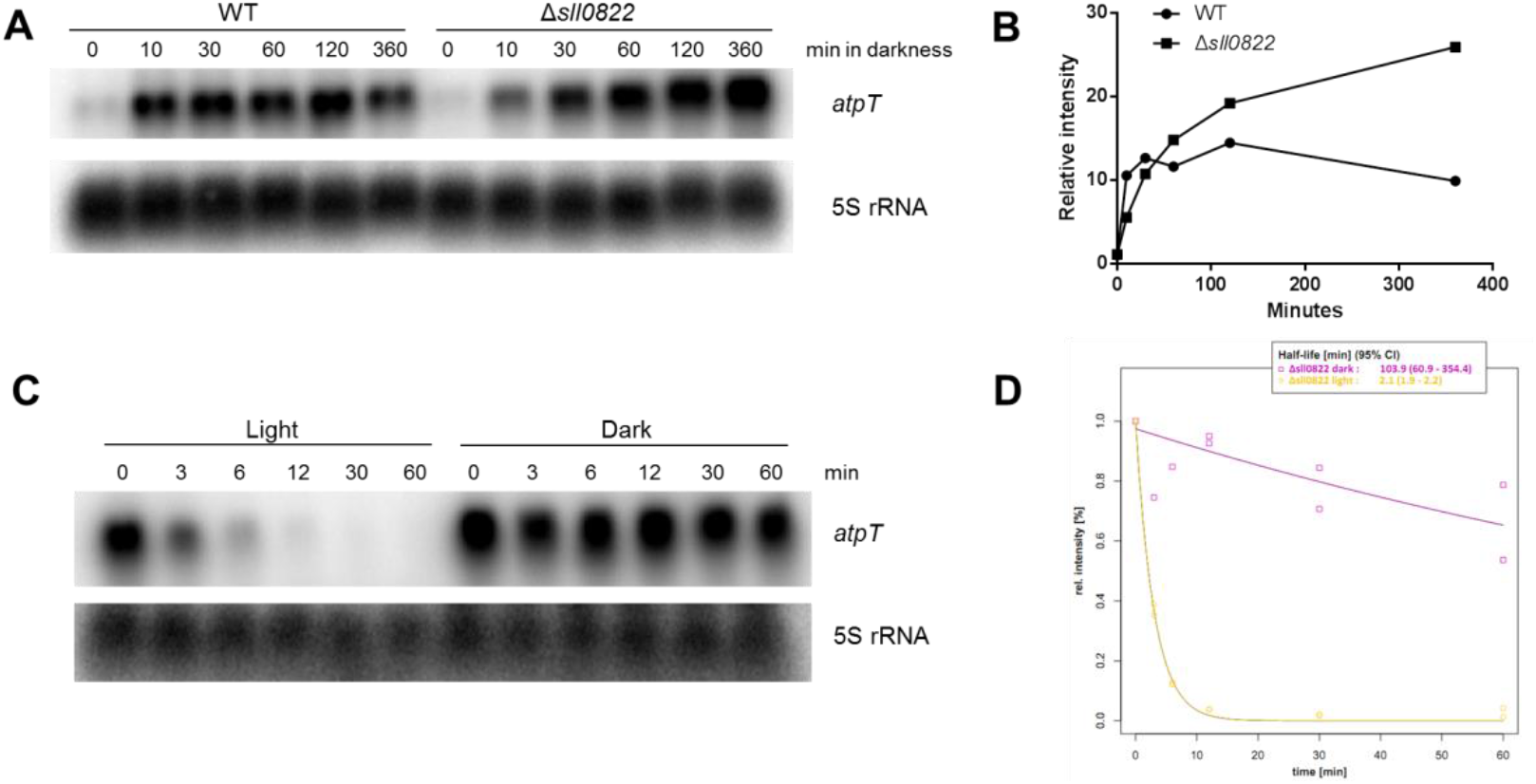
Accumulation and stability of the *atpT* transcript in the Δ*sll0822* mutant. (A) Northern blot analysis of the accumulation of the *atpT* transcript in both the wild type (WT) and Δ*sll0822* mutant. Cells with similar OD750 values were cultured under continuous light and then transferred to the dark. Samples were collected at the indicated time points. 5S rRNA was used as a loading control. (B) Levels of the *atpT* transcript in Panel (A) normalized to the respective 5S rRNA intensity plotted versus the time after rifampicin addition. (C) Northern blot analysis of the stability of the *atpT* transcript in the Δ*sll0822* mutant cultivated under either light or dark conditions. (D) Quantification of *atpT* transcript levels in Panel (C) and calculation of the decay curves as described in Fig. 3F.

The binding capacities of the two cyAbrBs were also tested under different pH and redox conditions that mimic different *in vivo* conditions. The cytosolic pH of the cyanobacterium *Synechococcus elongatus* PCC 7942 under light and dark conditions was reported to be 8.4 and 7.3, respectively (36). Here, a higher pH of 8.2 and a lower pH of 5 were tested. The binding capacity of cyAbrB1 to both the P_*atpT66*_-5’UTR and 5’UTR fragments was substantially decreased at pH 5. Only a slight shift was observed for P_*atpT66*_-5’UTR, and no shift was observed for the 5’UTR DNA fragment alone when high protein concentrations (16 μg) were added (Fig. 4C). The presence of DTT decreased the binding capacity of cyAbrB1 as well, while H_2_O_2_ exerted no discernible effect. In contrast, the binding of cyAbrB2 was not significantly altered by any of these conditions (Fig. S6).

### RpaB interacts with the *atpT* promoter *in vitro*

In the promoter deletion analysis we detected a dramatic effect if nucleotides in the AT-rich elements were substituted (Fig. 1). The selected region is similar to the HLR1 motif, the binding site for the RpaB transcription factor known to be involved in redox-dependent regulation of expression (20). Because this transcription factor wasn’t identified in the DNA affinity pull-down assay, we purified the His-tagged RpaB protein, and tested its binding with P_*atpT*_ promoter and its variant *in vitro* using EMSA. The results, as shown in Fig. 4E, did indicate RpaB binding to the P_*atpT66*_-5’UTR fragment *in vitro*, with a K_D_ value of 5.62E10-6. Moreover, unlike the two cyAbrB regulators, mutations within the P_*atpT*_ DNA fragment clearly affected the interaction with RpaB. This result suggests that RpaB is involved in the regulation of P_*atpT*_ promoter activity.

### Induction kinetics of *atpT* mRNA accumulation and stability are altered by the absence of cyAbrB2 *in vivo*

The interaction of the two cyAbrB proteins with the P_*atpT66*_ promoter was verified; however, convincing evidence for their involvement in *atpT* expression *in vivo* remains to be shown. Therefore, we studied deletion mutants. While cyAbrB1 is essential in *Synechocystis* 6803 (16), the accumulation of the *atpT* transcript in a cyAbrB2 knockout mutant can be investigated (Δ*sll0822*; (16)). The accumulation of the *atpT* transcript in both wild-type and Δ*sll0822* cells was monitored for 6 h after transfer to darkness, and the relative transcript levels were normalized to the respective 5S rRNA controls (Fig. 5A and B). Different kinetics were observed for *atpT* mRNA accumulation in Δ*sll0822* compared to wild type. In both strains, mRNA accumulation started within 10 min after transfer to the dark. However, the *atpT* transcript initially accumulated faster in wild-type cells than in Δ*sll0822*, while the *atpT* mRNA level gradually increased in Δ*sll0822*, finally reaching a higher level than in the wild type strain. This observation indicates that cyAbrB2 might be involved in the initial phase of *atpT* transcription or transcript accumulation after transfer to the dark.

Because we measured substantial differences in *atpT* mRNA stability under different conditions (Fig. 3), we tested the stability of the *atpT* transcript in Δ*sll0822* under both light and dark conditions (Fig. 5C and D). Compared with the *atpT* transcript stability in wild-type cells (Fig. 3), similar decay kinetics were observed in the light but a different pattern was observed under dark conditions in the Δ*sll0822* mutant. After transfer to darkness, the *atpT* transcript was degraded more slowly in Δ*sll0822* than in the wild-type strain (half-lives of 103.9 min and 33.2 min, 95% CI of 60.9-354.4 and 22.9-59.8, respectively). Therefore, cyAbrB2 plays some, possibly circumstantial, role in the degradation of the *atpT* mRNA in the dark.

## Discussion

### *atpT* promoter architecture and interacting proteins

AtpΘ is a recently reported ATP synthase inhibitor in cyanobacteria that accumulates under low-energy conditions and inhibits the futile ATP hydrolysis activity of ATP synthase (14). In the current study, the regulation of the *atpT* gene was studied in *Synechocystis* 6803, leading to the proposed model shown in Fig. 6. The activity of the *atpT* promoter was characterized in GFP-based reporter assays, which revealed that the combination of the *atpT* promoter and 5’UTR was necessary for maximum expression and activation under dark conditions (Fig. 1). Two AT-rich regions located between -45 and -60 relative to the TSS were essential for promoter activity, suggesting that this element is a potential transcription factor binding site. We noticed that this region is similar to the HLR1 motif, the binding site for the redox-responsive OmpR-type transcription factor RpaB (20). HLR1 motifs consist of two direct repeats, (G/T)TTACA(T/A)(T/A), separated by 2 nt. Its location relative to the TSS is compatible with an activating function of RpaB under low light conditions (20), which would be consistent with the observed regulation. RpaB was not enriched in the DNA immunoprecipitation experiment therefore we tested it separately. The found K_D_ values with the native promoter sequence were very similar to those of the strongly binding transcription factor cyAbrB2 while they were one order of magnitude lower with the mutated fragment (Fig. 4E) containing the same nucleotide substitutions that led to the loss of promoter function in the reporter assay (Fig.2F). Hence, this result strongly suggests that RpaB is involved in the regulation of *atpT* transcription.

**FIG 6.**
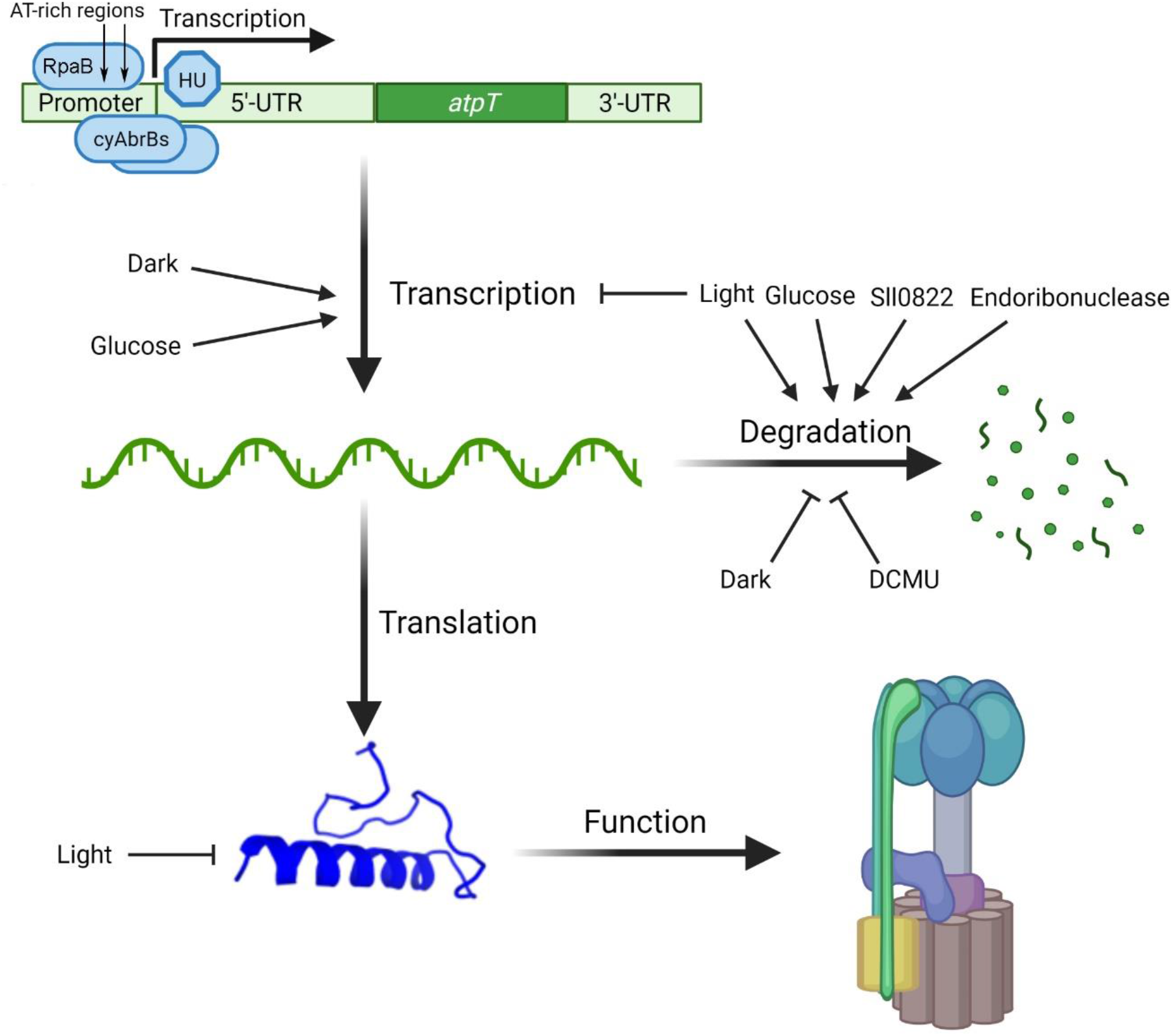
Proposed scenario for the regulation of *atpT* expression. Both the regulation of transcription and mRNA stability contribute to the *atpT* expression level. Two cyAbrBs bind to the promoter and 5’UTR of *atpT*, while the histone-like protein HU interacted mainly with the 5’UTR. Two AT-rich regions were identified in the *atpT* promoter that are relevant for regulation. These constitute HLR1 elements and bind RpaB. Multiple factors affect *atpT* expression, most prominently light/dark conditions or inhibition of electron transport by DCMU. The addition of glucose exerted two divergent effects: (i) a likely nonspecific stimulation of transcription and (ii) a substantially reduced *atpT* mRNA stability. The *atpT* gene product AtpΘ ultimately inhibits the ATP hydrolysis activity (reverse reaction) of F_0_F_1_ ATP synthase. As a third level of regulation, AtpΘ protein stability is dramatically reduced if cultures are transferred from the dark to light (14). The figure was drawn using BioRender (https://biorender.com/).

Instead, we identified one and two proteins that were enriched by binding to the *atpT* 5’UTR and promoter DNA fragments, respectively (Fig. 6). The protein Sll1712, interacting with the *atpT* 5’UTR fragment, is a conserved nucleoid-associated, or HU-like, protein. These basic, histone-like proteins bind DNA strongly and have an architectural role, but they have been implicated in transcriptional regulation via DNA looping, a mechanism that was documented to be present in cyanobacteria (37). The two proteins enriched with the *atpT* promoter fragment, cyAbrB1 and cyAbrB2, belong to the family of AbrB proteins that are strongly DNA-binding as well (38). All three proteins are small in size, abundant, and positively charged transcriptional regulators (the pI and molecular mass in kDa of cyAbrB1, cyAbrB2 and Sll1712 is 9.18/17.2; 8.63/13.9 and 8.69/10.8, respectively). Thus, also cyAbrB1 and cyAbrB2 possess some similarities with nucleoid proteins. AbrB-type DNA-binding proteins were characterized to bind to degenerate A/T-rich sequence elements, and their binding may lead to DNA bending (39). These characteristics are compatible with the identification of cyAbrB1 and cyAbrB2 here. Although cyAbrB2 was also reported to bind nonspecifically to DNA (16), we found that both cyAbrBs showed stronger binding capacity to the *atpT* promoter + 5’UTR than to the 5’UTR alone when lower protein-to-DNA ratios were used (Fig. 4A and B). Thus, the combination of the promoter and 5’UTR of *atpT* forms special structures that bind cyAbrBs. The DNA-binding activity of cyAbrB1 was diminished in the presence of DTT or H_2_O_2_ and affected by pH (Fig. 4C), suggesting that redox signals affect the *atpT* promoter to some extent *in vivo*. Although the structure of AbrB regulators from *Bacillus* has been studied (40), the pH-dependent regulation of their DNA binding capacity is still unclear.

The involvement of cyAbrBs in *atpT* expression is consistent with the previously observed dysregulation of *atpT* in the transcriptome-wide analysis of cyAbrB2 functions in the acclimation to low carbon (35). In that study, *atpT* (called *norf1* there) was assigned to the core set of upregulated protein-coding genes upon a 24 h shift from high (5% CO_2_) to low (24 h ambient air) inorganic carbon supply in the wild type strain. An upregulation of *atpT* was also observed in Δ*sll0822;* however, the mRNA levels were lower than those in the wild-type strain in the presence of both high and low inorganic carbon concentrations (Fig. S7). Consistent with this previous result, we observed different kinetics for *atpT* mRNA accumulation in light/dark-shifted Δ*sll0822* cells compared to wild-type cells (Fig. 5). These results clearly indicate a role for cyAbrB2 in sustaining the correct regulation of *atpT* transcription and that further regulators must be involved. One such regulator would be RpaB according to the results of this work.

### Conditionally different transcript stabilities are the major effector of *atpT* expression

Because the rapid and strong regulation of *atpT* expression during light/dark shifts cannot be explained by the promoter alone, the transcript stabilities under different conditions were further investigated. Under dark conditions, the addition of glucose led to the almost complete disappearance of the *atpT* transcript (Fig. 2A), consistent with previous analyses (14). Because we observed the opposite regulation of *atpT* promoter activity in the GFP reporter approach (Fig. 2C), we concluded that the reduced transcript stability under dark + glucose conditions (Fig. 3C) led to the observed disappearance of the *atpT* mRNA.

A similar observation was recorded under light conditions: the *atpT* promoter was active at a substantial level (Fig. 1C), while rapid transcript degradation (Fig. 3B) led to the observed low transcript accumulation (Fig. 2A). These posttranscriptional effects were clearly transcript-specific: while the *atpT* mRNA was rapidly degraded in the dark in the presence of glucose and even more rapidly in the light (Fig. 2F), a similar result was not observed for the *gfp* mRNA. Thus, a reasonable speculation is that the presence of glucose stimulated respiratory ATP production, triggering a posttranscriptional mechanism that prevented high AtpΘ levels. Another major effector was DCMU, which led to drastically prolonged *atpT* mRNA stabilities. DCMU interrupts the photosynthetic electron transport chain by blocking the Q_B_ plastoquinone binding site of photosystem II (PSII) (41). Therefore, the associated effects provide evidence that redox signaling is involved in this process. However, the effects of DCMU observed in the dark are unlikely to be related to PSII. DCMU was also shown to bind the cytochrome b_6_f complex in the binding site for plastoquinone in yeast (42). The cytochrome b_6_f complex in cyanobacteria is an integral part of the photosynthetic and respiratory electron transfer chain and is therefore relevant for dark respiratory electron transport. Instead, high DCMU concentrations, such those used here, were indeed reported to lower the dark respiration rate of the cyanobacterium *Anabaena halophytica* (43). Therefore, the application of DCMU in the dark certainly exerted a redox effect as well, probably by inhibiting the cytochrome b_6_f complex. Again, the effect of DCMU under both light and dark conditions was most pronounced on the *atpT* mRNA level. We conclude that the redox-mediated control of *atpT* mRNA stability is a major factor regulating AtpΘ expression.

Overall, our results show that the *atpT* promoter and *atpT* transcript accumulation are affected by multiple factors in addition to light or energy availability (Fig. 6). Because the native *atpT* 5’UTR was fused to the sf*gfp* coding sequence, the effect on mRNA stability must have been related to the *atpT* coding sequence or the 3’UTR. We noticed that putative endoribonuclease sites were identified at the beginning of the *atpT* coding sequence in a recent transcriptome-wide mapping study (44). The two largest of these endoribonuclease sites were mapped 5 and 22 nt downstream of the start codon; hence, cleavage at these sites would be consistent with the immediate inactivation of the *atpT* mRNA. Those sites were mapped in cells cultivated in light (a condition where the *atpT* mRNA is very short-lived) and likely are key sites for the destabilization of this mRNA. Following endoribonucleolytic cleavage, the transcript is likely rapidly degraded further by exonucleases. This endoribonuclease targeting the *atpT* mRNA is likely not RNase E because those sites became prominent upon the transient inactivation of RNase E by a temperature shift (44). A candidate is RNase J, especially since *Synechocystis* RNase J exhibited similar processing activities *in vitro* (45).

Collectively, our results present a striking parallel to observations of the regulation of eukaryotic IF1 inhibitors, which are less controlled at transcriptional level as well, although there the regulation appears to occur mainly at the level of IF1 translation or protein stability and not mRNA stability (46). The elucidation of mechanisms involved in the function and regulation of putative homologs and analogs in plants and in cyanobacteria is an interesting topic for further research.

## Materials and Methods

### Strain construction and cultivation

The Δ*sll0822* (Δ*cyabrB2*) mutant (16) was provided by Y. Hihara, Saitama University, Japan. *Synechocystis* 6803 was cultured in BG11 medium buffered with 20 mM TES (pH 8.0) (17) with gentle agitation or on solid agar (1.5% Kobe I agar) plates under constant illumination of approximately 40 µmol photons m^-2^ s^-1^ at 30 °C. If appropriate, antibiotics were added (5 μg ml^-1^ gentamicin for GFP reporter strains, 10 μg ml^-1^ kanamycin for the Δ*sll0822* mutant). For incubations in the dark, flasks were wrapped with tin foil. If indicated, 1% glucose [w/v] and/or 3-(3,4-dichlorophenyl)-1,1-dimethylurea (DCMU), carbonyl cyanide m-chlorophenylhydrazone (CCCP), or methyl viologen were added at concentrations of 50, 10 or 6 μM. Rifampicin was added at 300 μg/ml from a 50 mg/ml stock solution in dimethyl sulfoxide to study RNA stability.

*E. coli* TOP10 was used for plasmid construction, *E. coli* BL21 (DE3) for protein expression, and *E. coli* J53/RP4 as a helper strain for conjugation. *E. coli* cells were grown in liquid LB medium (10 g/l bacto-tryptone, 5 g/l bacto-yeast extract, and 10 g/l NaCl) with continuous agitation or on 1.5% [w/v] Bacto agar plates at 37 °C. If required, 5 μg ml^-1^ gentamicin, 50 μg ml^-1^ ampicillin or 50 μg ml^-1^ kanamycin was added.

For GFP assays, the plasmids Pgfpconstruct_1, 3, 4, 5 and 6 were constructed through AQUA cloning (18) by fusing the promoter, sf*gfp* and pVZ322 backbone fragments amplified by PCR. The plasmid Pgfpconstruct_2 was obtained using AQUA cloning based on the plasmid Pgfpconstruct_1, plasmids Pgfpconstruct_mut1, 2, 3 and 5 were constructed using AQUA cloning based on the plasmid Pgfpconstruct_3, and plasmid Pgfpconstruct_mut4 was constructed using AQUA cloning based on the plasmid Pgfpconstruct_mut3. For the expression of recombinant cyAbrB1 and cyAbrB2, the plasmids pNHis-0359 and pNHis-0822 were constructed through AQUA cloning and transformed into *E. coli* BL21 (DE3). All primers are listed in Table S1, and all strains are listed in Table S2.

### Recombinant protein purification

For the overexpression His-tagged cyAbrB1 and cyAbrB2, ten milliliters of *E. coli* BL21 (DE3) overnight cultures containing pNHis-0822 or pNHis-0359 were inoculated into 1 L of LB medium supplemented with ampicillin and cultured at 37 °C with continuous agitation. The expression of recombinant proteins was induced by adding isopropyl β-d-1-thiogalactopyranoside (IPTG) to a concentration of 0.5 mM when the OD600 reached 0.6. After 4 h, the cells were collected by centrifugation at 3,000 *g* for 10 min. Pellets were resuspended in PBS (137 mM NaCl, 2.7 mM KCl, 10 mM Na_2_HPO_4_, and 1.8 mM KH_2_PO_4_, pH 7.5) supplemented with Protease Inhibitor Cocktail and then broken using a cell disruptor (Constant Systems Limited). The lysates were centrifuged at 12,000 *g* for 30 min, and the supernatants filtered through a 0.45 μm filter. The recombinant proteins were then purified using a 1 ml HisTrap HP column and ÄKTA™ system (GE Healthcare) according to the manufacturer’s instructions. For the overexpression of His-tagged RpaB, a similar procedure was carried out but using *E. coli* BL21 (DE3) pLysS as described previously (19, 20).

### GFP assays

GFP fluorescence in *Synechocystis* 6803 was measured using a VICTOR3 multiplate reader (PerkinElmer). The cells were cultured to an OD750 of 0.5, and 200 μl aliquots were transferred to a 96-well microplate. The fluorescence was measured with an excitation filter F485, emission filter F535, a measurement time of 1.0 s, and a CW-lamp energy 21673. In parallel, absorbance was measured at 750 nm (CW-lamp filter 750/8 nm, measurement time 1.0 s) and then used to normalize the GFP intensity. Since the adjusted GFP intensity decreased with the increase in OD750 (Fig. S1), probably due to self-shading, the cultures were adjusted to equal OD750 values the day before the measurement.

### Protein extraction, Western blotting and polyacrylamide gel electrophoresis

A 600 ml culture was cultured in a 2 L flask until reaching an OD750 of 0.8 and then divided into 50 ml aliquots to analyze the GFP level in the GFPconstruct_1 strain cultured under different conditions. One aliquot was collected as a light culture control, while the others were treated as indicated. *Synechocystis* 6803 cells were harvested by centrifugation (4,000 *g*, 10 min, 4 °C), resuspended in PBS containing a protease inhibitor cocktail, and disrupted mechanically using a Precellys 24 homogenizer (Bertin Technologies). Debris was removed by centrifugation at 1,000 g for 1 min at 4 °C, and the protein concentration in the supernatant was measured using the Bradford assay. Before loading, protein samples were boiled with 1x protein loading buffer at 95 °C for 10 min.

For denaturing SDS–PAGE, samples were incubated with 1x loading buffer at 95 °C for 10 min and then separated on 15% Tris-glycine SDS PAGE or Tricine–SDS PAGE gels (21). For Western blotting, the gels were transferred to Hybond™-ECL membranes (Cytiva) using semidry electroblotting. Blotted membranes were blocked with 3% skim milk dissolved in TBST (20 mM Tris, pH 7.6, 150 mM NaCl, and 0.1% Tween-20) and then sequentially incubated with rabbit anti-GFP primary antisera (1:2000 dilution; Sigma–Aldrich) and secondary antisera (1:10,000 anti-rabbit antisera; Sigma–Aldrich). Signals were detected with ECL start Western blotting detection reagent (Cytiva) using a chemiluminescence imager system (Fusion SL, Vilber Lourmat). Western blots were repeated twice independently.

Native SDS–PAGE (NSDS–PAGE) was conducted as described previously (22) with modifications. Protein samples were mixed with 1/3 volume of 4 x loading buffer (100 mM Tris-HCl, 0.01875% Coomassie G-250, and 10% glycerol, pH 8.5) and then loaded on a native glycine polyacrylamide gel (same recipe as the 15% Tris-glycine SDS gel except for the omission of SDS). Electrophoresis was performed in NSDS– PAGE running buffer (50 mM Tris-HCl, 200 mM glycine, and 0.0375% SDS, pH 7.3) at 4 °C.

### RNA isolation and Northern blotting

*Synechocystis* 6803 cells were harvested by vacuum filtration on hydrophilic polyethersulfone filters (Pall Supor®-800, 0.8 µm), followed by snap freezing in liquid nitrogen; the entire collection procedure took approximately 30 s. Isolation of total RNA and Northern blotting procedures were performed as described previously (14). Primer and oligonucleotide sequences are listed in Table S1. Signals were visualized using Typhoon FLA 9500 (GE Healthcare) and Quantity One software (Bio-Rad). Quantity One software was also used to quantify *atpT* and 5S rRNA transcript levels. For subsequent normalization, the *atpT*/5S rRNA ratio at time 0 under each condition was set to 1, and the values at the following time points were calculated accordingly. All Northern blots were independently performed at least twice.

### Calculation of the transcript half-life based on Northern hybridization results

The stability of the *atpT* mRNA was measured after adding rifampicin and transferring the cultures to the indicated conditions simultaneously. RNA samples prepared before and at the indicated times after rifampicin addition were analyzed using Northern hybridization. Hybridization results were quantified using Quantity One® (Bio-Rad) software, and 5S rRNA was used as a loading reference. Relative intensities of each blot were normalized to the relative intensity before rifampicin addition (normalized relative intensity of 1 for time 0). The normalized relative intensities of *atpT* transcripts were fitted to the exponential decay function *I*(*t*) = *I*(*t*_0_)*e*^−*λt*^ with the R nls2 function using start values in the range from 0.001 to 0.5 for the decay constant λ and from 0.2 to 2 for the start intensity *I*(*t*_0_). The half-life was calculated using the equation 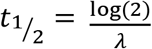.

### Protein pull-down and mass spectrometry

DNA affinity pull-down assays using P_*atpT66*_-5’UTR and the *atpT* 5’UTR DNA fragments were performed as described previously with modifications (23). Three liter cultures of *Synechocystis* 6803 were collected by centrifugation at 7,000 *g* for 10 min, washed twice with PBS (pH 7.5), and resuspended in 4 ml of BS/THES buffer (0.5 mM MgCl_2_, 50 μM ZnSO_4_, 50 μM FeCl_2_, 50 μM MnCl_2_, and 50 μM EDTA) supplemented with Protease Inhibitor Cocktail. The cells were disrupted in a Precellys 24 homogenizer, and debris was pelleted by centrifugation at 13,000 *g* for 30 min at 4 °C. Six hundred microliters of the resulting supernatant were first incubated with 200 μl of uncharged Dynabeads M-280 Streptavidin (Thermo Fisher Scientific) for 20 min to minimize nonspecific binding before an incubation with beads loaded with the bait for 1 h at room temperature. Before the incubation with lysate, Dynabeads were washed and prepared according to the manufacturer’s instructions. The biotinylated P_*atpT66*_-5’UTR and the *atpT* 5’-UTR DNA fragments were amplified by PCR using the primers listed in Table S1 and purified with the Macherey-Nagel™ NucleoSpin™ Gel and PCR Clean-up Kit.

After the incubation with cell lysate, beads charged with baits and bound proteins were washed with BS/THES buffer 5 times, and proteins were eluted with elution buffer (25 mM Tris–HCl and 250 mM NaCl). Protein elutes were precipitated overnight at 4 °C by adding trichloroacetic acid to a final concentration of 15%. Proteins were pelleted by centrifugation at 13,000 *g* for 30 min at 4 °C and then analyzed on a 16%/6 M urea Tricine-SDS gel. The gel was stained with InstantBlue (Expedeon), and eventually, samples were prepared and analyzed using mass spectrometry as previously described (14). The proteomics data have been deposited at the ProteomeXchange Consortium via the PRIDE partner repository (24) with the dataset identifier PXD027801.

### Electrophoretic mobility shift assay and calculation of K_D_ value

Cy3-labeled probes were amplified by PCR using the primers listed in Table S1 and purified with the Macherey-Nagel™ NucleoSpin™ Gel and PCR Clean-up Kit. Reaction conditions for binding protein to DNA were 12 mM HEPES, 4 mM Tris, 12 mM KCl, 1 mM EDTA, 1 mM EGTA, 12% glycerol, and 1 μg of LightShift Poly (dIdC) (Thermo Fisher Scientific). Samples were run on 3% agarose gels in 0.5x TBE (45 mM Tris, 45 mM boric acid, and 1 mM EDTA, pH 8.0) and visualized using Typhoon FLA 9500 (GE Healthcare) and Quantity One® software (Bio-Rad).

The equilibrium dissociation constant (K_D_) between DNA probe and protein was calculated based on the intermediate lanes of the EMSA assays. K_D_ was calculated as 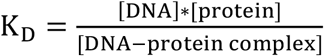, where the concentration of DNA-protein complex was calculated according to the relative signal intensity versus the control lane without proteins). When available, two K_D_ values were calculated for each DNA-protein pair, and the mean was used as the final K_D_ value. Higher K_D_ value indicates lower affinity between the DNA probe and the protein.

### Statistical analysis

Statistical analyses were performed with GraphPad Prism 6.0 software (GraphPad Software, Inc., San Diego, CA). Unpaired two-tailed *t* tests were performed, as shown in Tables S5 and S6. Differences between groups were considered significant at a *P* value of <0.05 and highly significant at a *P* value of <0.01.

## Data availability

The datasets produced in this study are available in the following databases:

- Mass spectrometry raw data were deposited at the ProteomeXchange Consortium (http://proteomecentral.proteomexchange.org) via the PRIDE partner repository (24) under the identifier PXD027801.

## SUPPLEMENTAL MATERIAL

Supplemental material is available online only.

Fig. S1 to Fig S7.

Tables S1 to S6.

## Acknowledgments

We are grateful for the gift of the Δ*sll0822* mutant lacking cyAbrB2 by Y. Hihara, Saitama University, Japan. This study was funded by the German Research Foundation (DFG) priority program SPP2002 “Small Proteins in Prokaryotes, an Unexplored World” (grant HE 2544/12-2 to WRH, grant HA 2002/22-2 to MH and grant BE 3869/5-2 to DöB) by the DFG-funded research training group BioInMe 322977937/GRK2344 to K.S. and W.R.H., and by a China Scholarship Council grant to K.S.

## Authors’ contributions

KS performed the molecular genetic and biochemical analyses in *Synechocystis* 6803, and DB and SM performed mass spectrometry analyses. MH provided scientific input for improving the experimental design and physiological interpretation. JG performed statistical analyses of RNA stability. WRH designed the study, and all authors analyzed the data. KS and WRH drafted the manuscript with input from all authors. All authors read and approved the final manuscript.

