## Supplemental Figures S1 to S7 for "Expression of the cyanobacterial F_0_F_1_ ATP synthase inhibitor AtpΘ depends on small basic DNA-binding proteins and differential mRNA stability"

### SUPPLEMENTAL MATERIAL

**Supplementary figures**

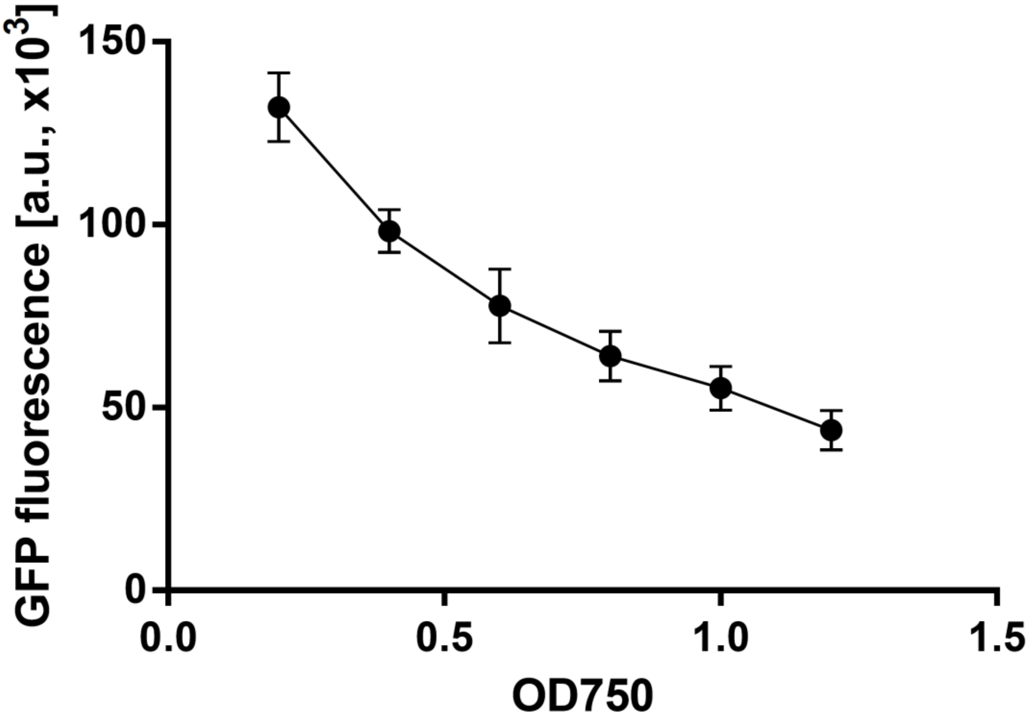


**FIG S1** The correlation between GFP intensity and OD750. The exponential phase culture of the strain containing GFPconstruct_1 (OD750 approximately 1.2) was diluted as indicated, and the GFP intensities were measured immediately afterward.

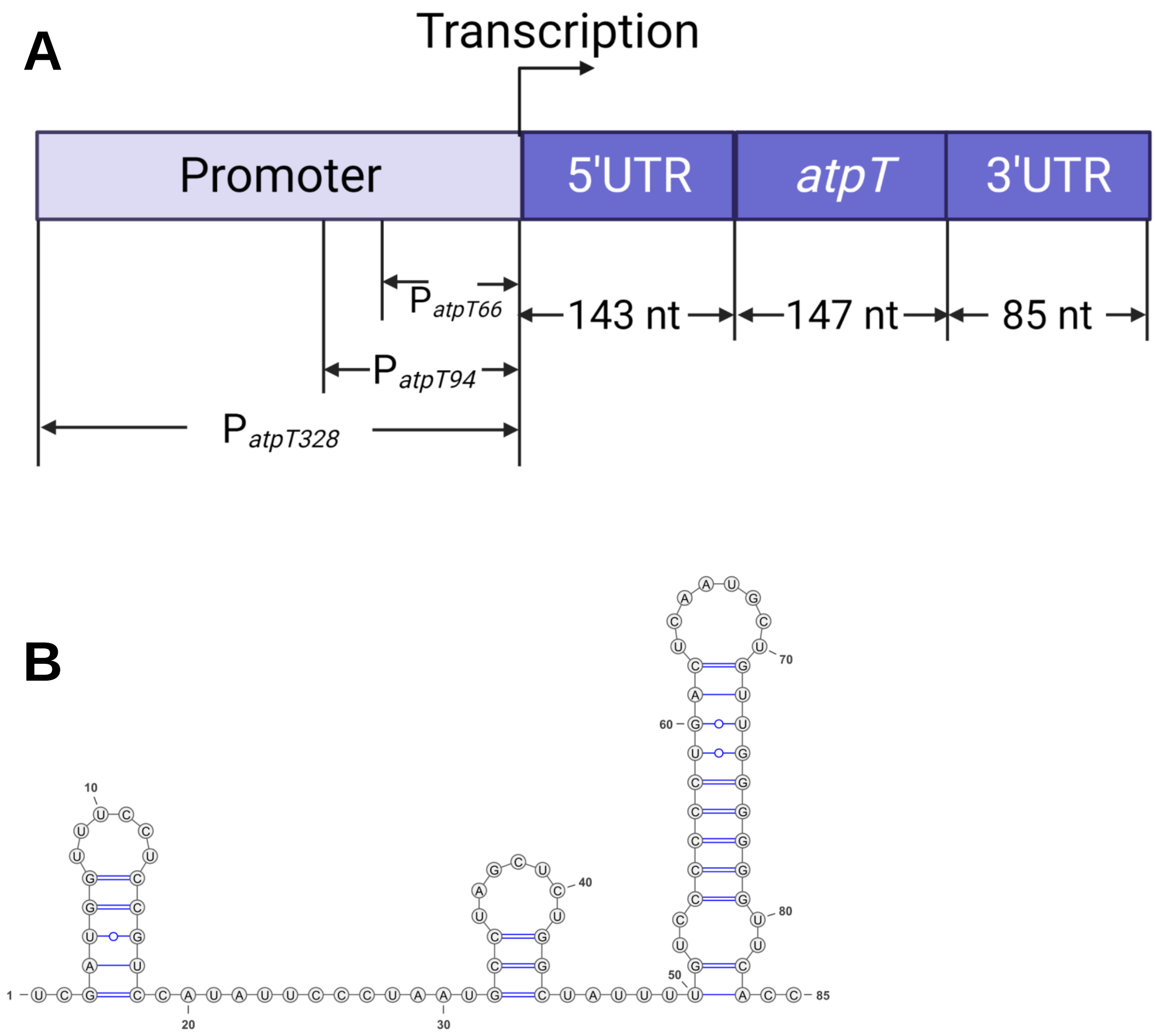


**FIG S2** Structure of the *atpT* gene and transcript in *Synechocystis* 6803. (A) The *atpT* transcript consists of a 143 nt 5’UTR, 147 nt coding sequence, and 85 nt 3’UTR. Three *atpT* promoter fragments encompassing 66 nt, 94 nt or 328 nt upstream of the TSS were investigated, called $P_{atpT66}$, $P_{atpT94}$ and $P_{atpT328}$, as indicated. (B) Predicted RNA secondary structure of the 3’UTR.

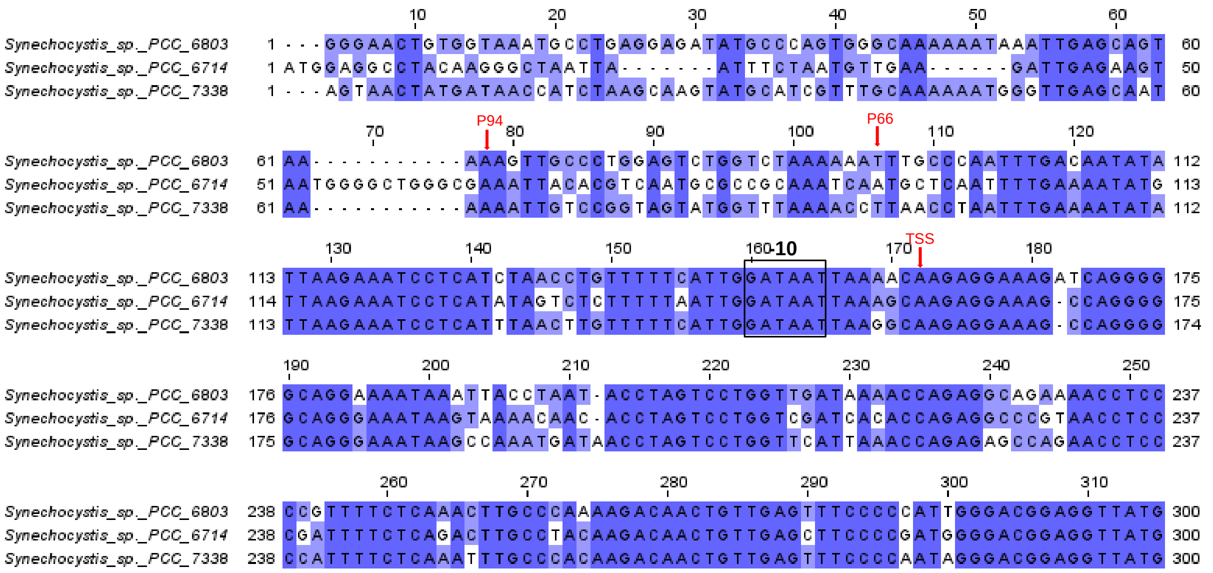

**FIG S3** Alignment of the promoter and 5'UTR regions of *atpT* homologs in three *Synechocystis* strains. Dark blue shading indicates identical residues in all three compared strains, and light blue shading indicates identity in the sequences of any two strains. Red arrows mark the TSSs identified previously for *Synechocystis* 6803 and *Synechocystis* 6714 (1, 2). The positions -66 and -94 relative to the TSS in *Synechocystis* 6803 are indicated. These fragments were chosen to define fragments for experimental analyses (compare Fig. S2a). The -10 element according to the mapped TSSs in strains 6803 and 6714 is boxed. For the alignment, 300 nt upstream of the respective *atpT* start codon was selected using MUSCLE (3). Notably, the coding sequence in all three strains starts with two consecutive ATG codons. Here, we considered the second codon as the authentic start codon, while the first was considered the last codon of the 5'UTR. The sequence data were derived from GenBank files NC_000911.1 (*Synechocystis* 6803), CP007542.1 (*Synechocystis* 6714), and CP054306.1 (*Synechocystis* 7338).

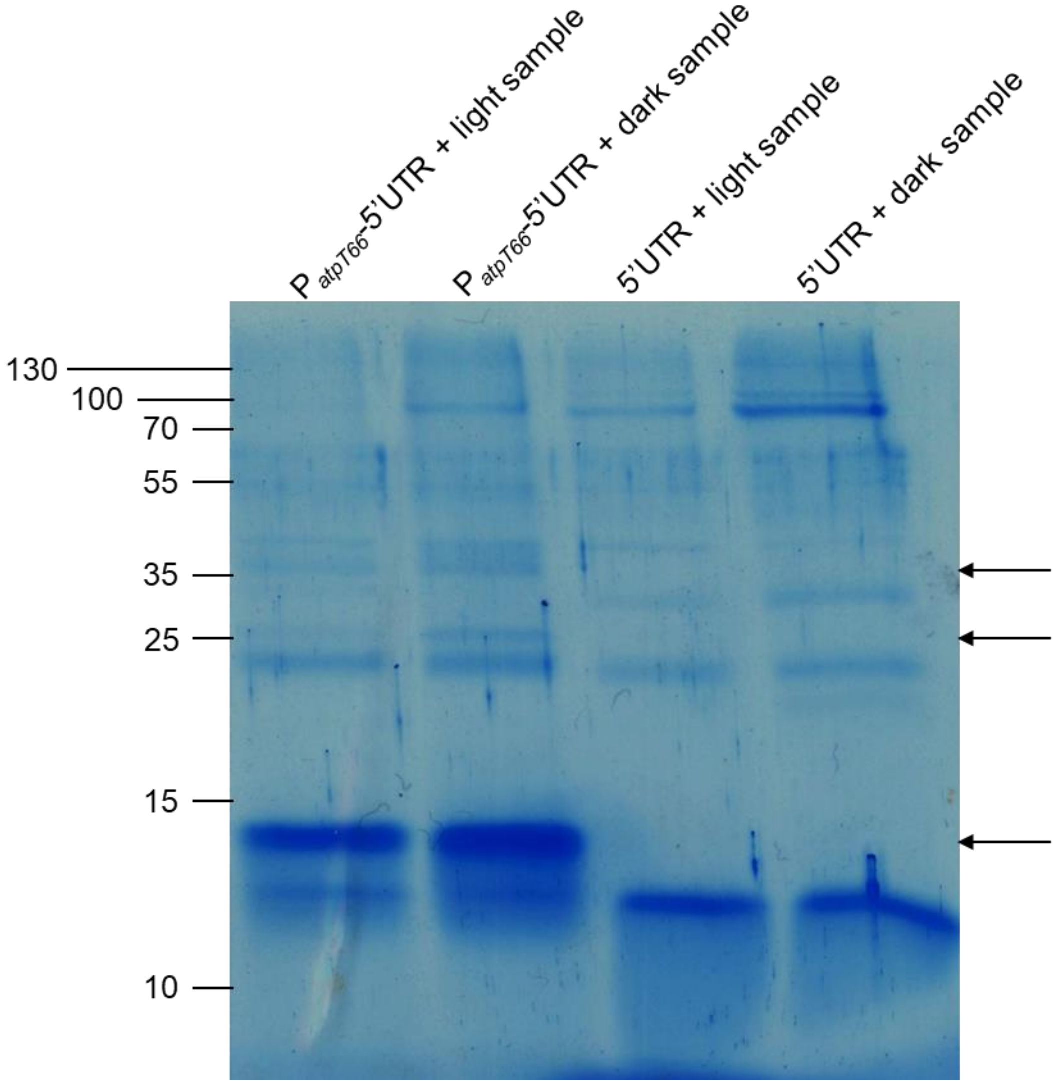


**FIG S4** Tricine SDS–PAGE gel showing the sample pulled down using different biotinylated DNA fragments. Two DNA fragments containing the 66 nt-long promoter variant and the 5'UTR ($P_{atpT66}$-5'UTR) or the *atpT* 5'-UTR were used. These two fragments were incubated with total protein samples isolated from wild type *Synechocystis* 6803 cultured under continuous light or after 12 h in the dark. The arrows indicate bands of significantly enriched proteins interacting with the $P_{atpT66}$-5'UTR or the *atpT* 5'-UTR DNA fragment. The same samples were subsequently subjected to mass spectrometry analysis. PageRuler™ Prestained Protein Ladder (Thermo Scientific, 10 to 180 kDa) was used as a molecular mass marker.

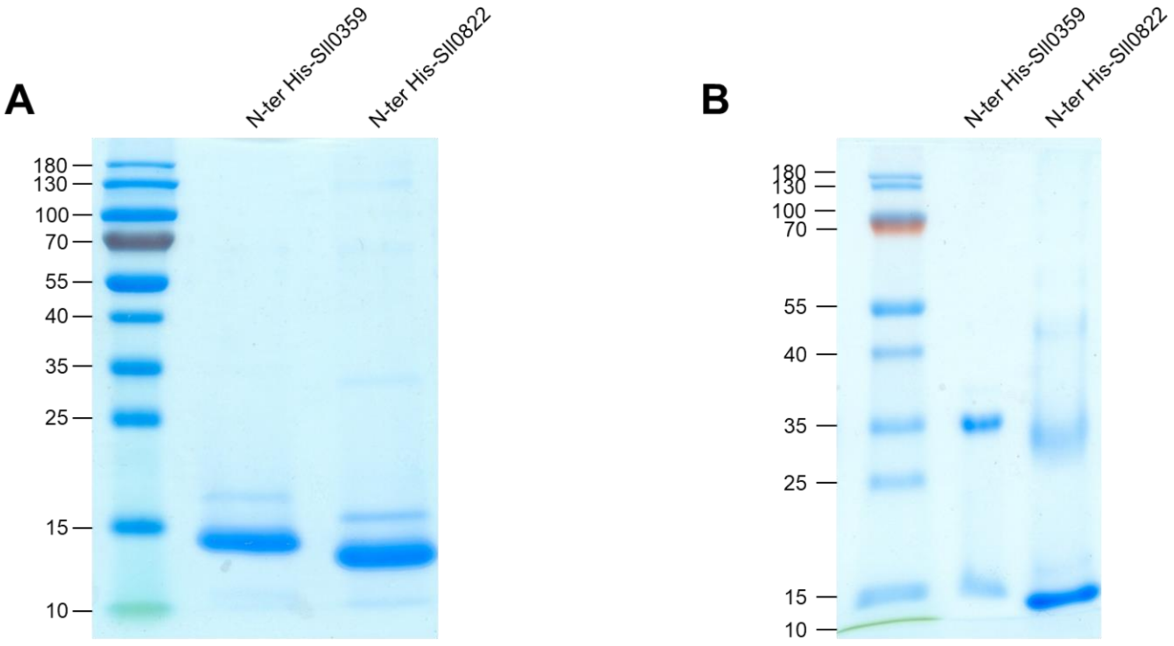


**FIG S5** PAGE analysis of the purified Sll0822 and Sll0359 proteins. The recombinant 6*His-tagged Sll0822 and Sll0359 proteins were purified from *E. coli* BL21 (DE3) cells, and 5 µg of each purified protein were loaded on (A) SDS–PAGE or (B) native SDS–PAGE gels for analysis.

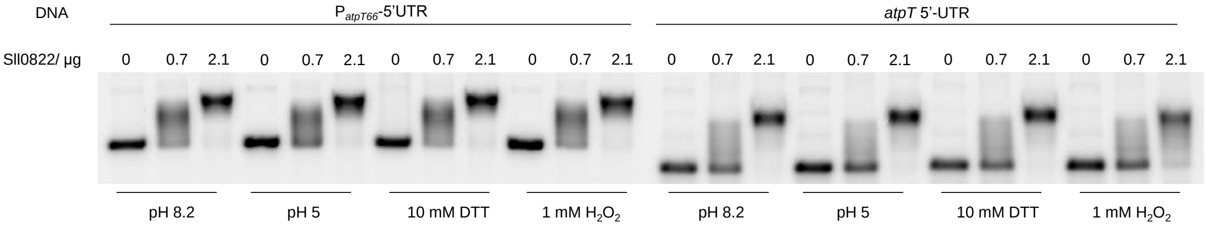


**FIG S6** Electrophoretic mobility shift assay. The binding of cyAbrB2 to the $P_{atpT66}$-5'UTR or *atpT* 5'UTR DNA fragments under different conditions was tested by performing gel shift assays.

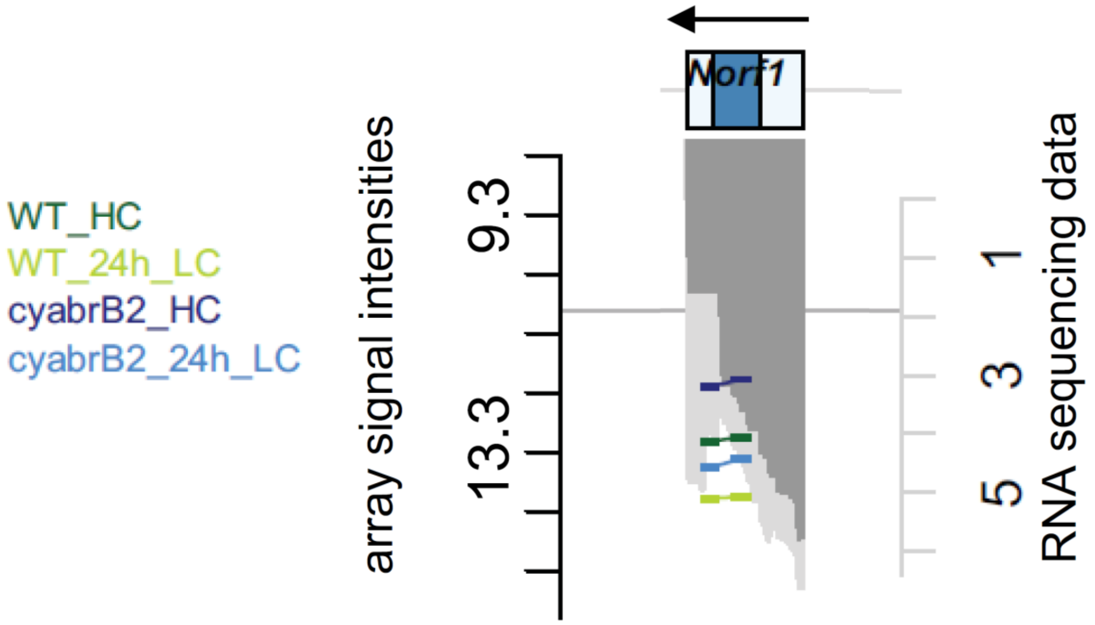


**FIG S7** Genomic locus of *atpT* in Orf *et al.* (4)(called *norf1* there) with mapped array probes (note that the gene is located on the reverse strand, the direction of transcription is indicated by the horizontal arrow). *Synechocystis* 6803 WT and the Δ*cyabrB2* mutant were compared in the presence of high (HC, 5% $CO_2$) and low (LC, 24 h ambient air) inorganic carbon supplies. The array signal intensities are presented as $log_2$ values and color-coded as indicated. The gray graphs represent RNA sequencing data presented as $log_2$ read numbers, which were extracted from Mitschke *et al.* (5).
